# Inferring Tumor Progression in Large Datasets

**DOI:** 10.1101/2020.06.18.159228

**Authors:** Mohammadreza Mohaghegh Neyshabouri, Seong-Hwan Jun, Jens Lagergren

## Abstract

Identification of mutations of the genes that give cancer a selective advantage is an important step towards research and clinical objectives. As such, there has been a growing interest in developing methods for identification of driver genes and their temporal order within a single patient (intra-tumor) as well as across a cohort of patients (inter-tumor). In this paper, we develop a probabilistic model for tumor progression, in which the driver genes are clustered into several ordered driver pathways. We develop an efficient inference algorithm that exhibits favorable scalability to the number of genes and samples compared to a previously introduced ILP-based method. Adopting a probabilistic approach also allows principled approaches to model selection and uncertainty quantification. Using a large set of experiments on synthetic datasets, we demonstrate our superior performance compared to the ILP-based method. We also analyze two biological datasets of colorectal and glioblastoma cancers. We emphasize that while the ILP-based method puts many seemingly passenger genes in the driver pathways, our algorithm keeps focused on truly driver genes and outputs more accurate models for cancer progression.

**Author summary:** Cancer is a disease caused by the accumulation of somatic mutations in the genome. This process is mainly driven by mutations in certain genes that give the harboring cells some selective advantage. The rather few driver genes are usually masked amongst an abundance of so-called passenger mutations. Identification of the driver genes and the temporal order in which the mutations occur is of great importance towards research and clinical objectives. In this paper, we introduce a probabilistic model for cancer progression and devise an efficient inference algorithm to train the model. We show that our method scales favorably to large datasets and provides superior performance compared to an ILP-based counterpart on a wide set of synthetic data simulations. Our Bayesian approach also allows for systematic model selection and confidence quantification procedures in contrast to the previous non-probabilistic progression models. We also study two large datasets on colorectal and glioblastoma cancers and validate our inferred model in comparison to the ILP-based method.

## Introduction

Tumor progression is caused by somatic evolution in which genes are randomly mutated and so-called driver mutations confer the tumor a selective advantage [1]. Several properties of somatic evolution of tumors have been studied intensively and are today more approachable than ever before, for instance,

1. What is the number of driver mutations in a tumor or the average across a collection?
2. Which genes acquire driver mutation, i.e., are so-called driver genes, in contrast to passenger genes?
3. What are the dependencies among driver mutations and, in particular, in which order do they occur?

Our main interest is in developing methods to resolve these questions as they have immense potential towards the identification of targets for new drugs and the development of patient-specific treatment plans.

The fundamental difficulty of answering these questions lies in the successful identification of driver mutations amongst an abundance of passenger mutations. Without prior information, there is no way to identify driver mutations in a single tumor. Over-representation of a gene among those mutated across a large tumor collection is, however, a useful driver gene identification approach. This approach is now feasible with the availability of mutation data through several large scale cancer sequencing efforts, e.g., The Cancer Genome Atlas (TCGA) as well as the International Cancer Genome Consortium (ICGC).

Cancer progression has been extensively studied through the application of various models capturing dependencies between mutated genes, such as oncogenetic trees [2–6] and conjunctive Bayesian networks [7, 8]. More complex models are also used to study the cancer progression [9–11]. However, these models are computationally hard to train, and hence, not applicable for big datasets composed of large numbers of samples and genes, and in the abundance of passenger mutations. On the other hand, studying the cancer progression using basic models at the gene level seems to be insufficient to fully model somatic evolution in cancer [12]. The effects of a driver gene mutation are often mediated by a biological pathway or a protein complex that the gene belongs to. Consequently, a set of genes associated with the same biological pathway or protein complex may affect a tumor in the very same way when mutated, and the selective advantage provided to the tumor by one may exhaust that of any other, which would make the genes be mutated in a mutually exclusive manner. These mechanisms can also provide a partial explanation for the observed heterogeneity of cancer mutations. The mutual exclusivity in driver genes has been studied using various models including [13, 14]. These methods have later been extended to the cancer progression context, where the aim is to identify mutually exclusive driver pathways and their temporal ordering, simultaneously. In [15], the objective is to learn a conjunctive Bayesian network with modules of mutually exclusive genes at the nodes of the network. However, the training procedure is a heuristic algorithm that suffers from computational complexity and scalability issues. In [16], the aim is to find a set of linearly ordered mutually exclusive driver pathways using integer linear programming (ILP). This method can potentially be applied in the presence of passenger mutations and on datasets composed of tens of genes and hundreds of patients.

In this paper, we develop a probabilistic model of mutually exclusive linearly ordered driver pathways. We design a sampling based inference algorithm to train our model. Using an extensive set of experiments we demonstrate our method’s superior performance and scalability to large datasets in comparison to the ILP-based algorithm in [16]. We also analyze two biological datasets on colorectal adenocarcinoma and glioblastoma and demonstrate our superior performance on these datasets in comparison to the ILP-based counterpart [16].

## Linear Progression Model

In this section, we start by illustrating linear pathway progression in cancer using the example model in Fig 1. We then introduce our probabilistic model for this process for which we subsequently develop sampling-based inference algorithms in the next section.

**Fig 1.**
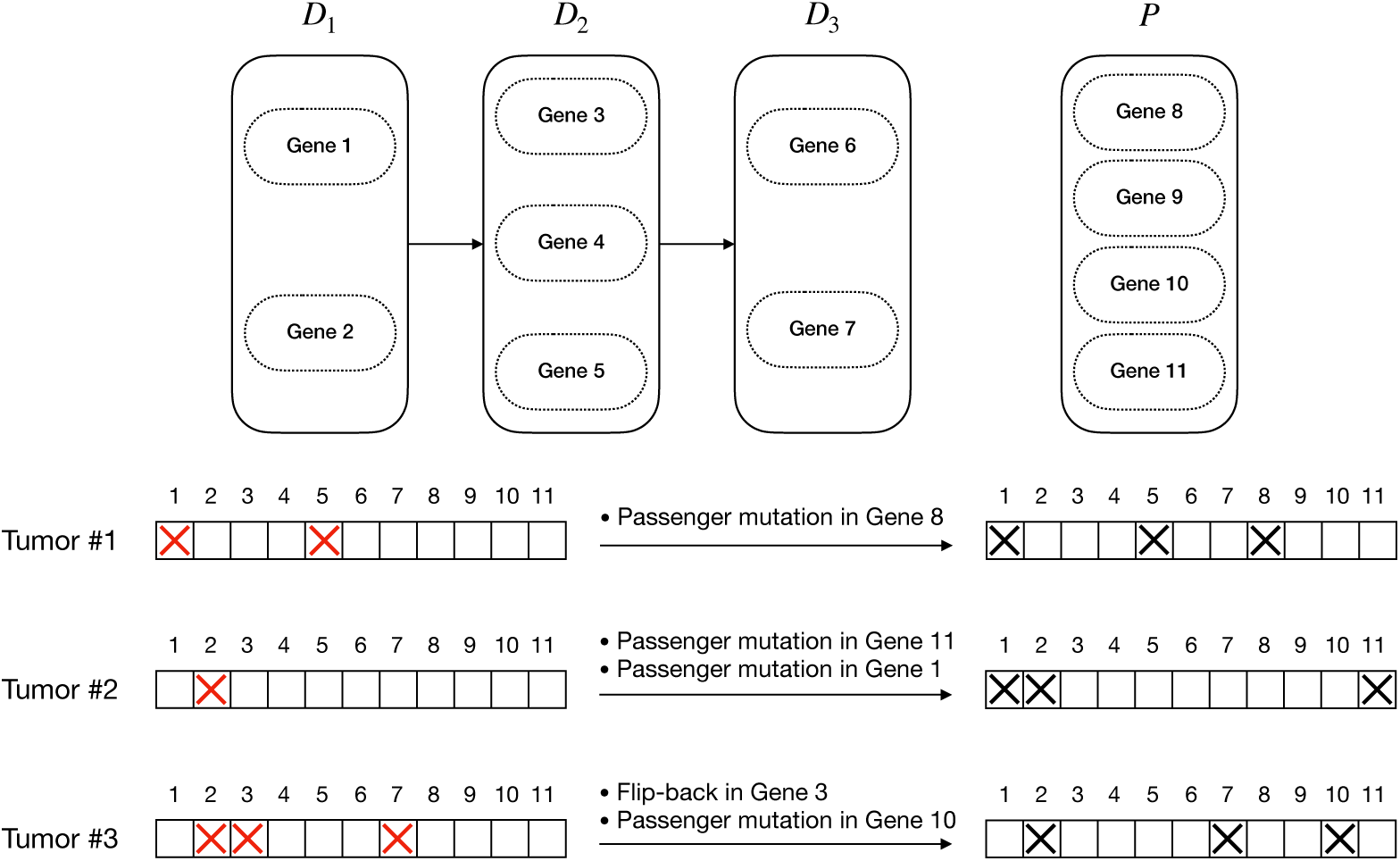
An example of a linear pathway progression model and three tumors generated following the model. The left-hand side gene status bars show the underlying mutational status of the genes with the red cross signs showing the driver mutations. The right-hand side gene status bars show the observed mutation status of genes, where passenger mutations and flip-back events make it hard to infer the true progression model from the observations.

### An illustration of the model

A linear pathway progression model of a specific cancer type (or sub-type) is defined as an ordered set of several sets of driver genes. We call these sets of genes *driver pathways*. We refer to the set of genes not included in the driver pathways as the *set of passengers*. Based on this model, cancer starts with a mutation in one of the genes in the first driver pathway. This mutation provides the harboring cells with some selective advantages and the tumor progresses to stage 1. As time goes on, the tumor may progress to stage 2 by acquiring some mutation in one of the genes in the second pathway adding more selective advantage to the tumor. The tumor can continue its progression further in the same way.

Our goal is to infer the driver pathways using a set of binary vectors showing the observed mutation status of a set of genes in various tumors. However, this task is not as simple as it seems to be. The vast majority of genes (belonging to the set of passengers) do not play an important role in the progression of the tumor. Hence, even though they may get mutated, these mutations are passenger mutations. Moreover, according to this model, if a pathway is already mutated due to a mutation in one of its constituting genes, no mutation in the other genes in the pathway can give the cells any more selective advantage. Therefore, such mutations will be considered as passenger mutations as well. Working with bulk data, we have a preprocessing step outputting the binary mutation status for each gene (1 if mutated, 0 otherwise). As a result, some actual driver mutations may get lost because of their small cellular prevalence for instance. We refer to these kinds of errors as flip-back events. For example, consider the progression model in Fig 1. As shown in the left-hand side mutation status vectors, tumors 1, 2, and 3 have the cancer stages of 2, 1, and 3, respectively. However, the passenger mutations and flip-back events lead to our observations shown in the right-hand side mutation status vectors.

#### Algorithm 1

Generative process

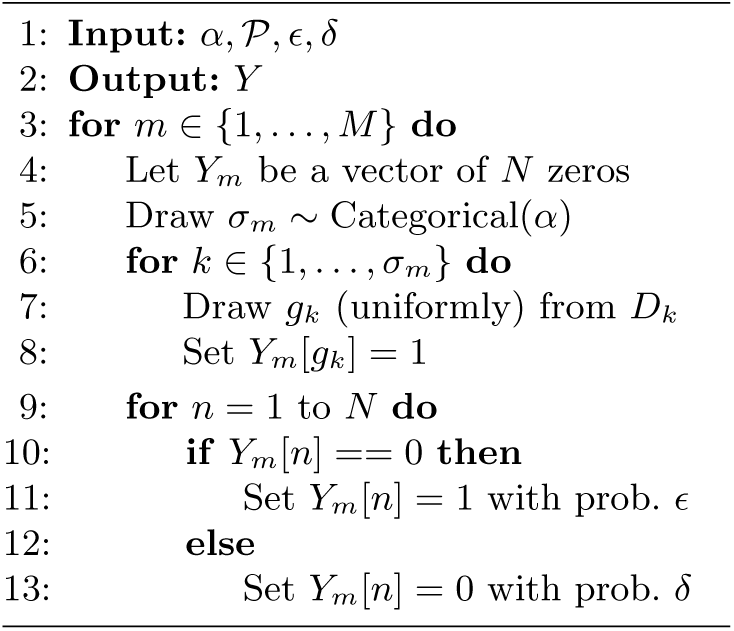

### Notation

Consider a set of *N* genes indexed from 1 to *N*. Let 𝒫 = (*D*_1_, *D*_2_, …, *D*_*L*_, *P*) be an ordered partition of the set of indices {1, …, *N*}. If each gene index is assigned to exactly one of the sets in 𝒫 and *D*_1_ to *D*_*L*_ are not empty, then 𝒫 is a *linear pathway progression model* of length *L*. We refer to *D*_1_, …, *D*_*L*_ as the driver pathways and *P* as the set of passengers.

The observed data consist of a mutation matrix *Y* ∈ {0, 1}^*M×N*^ for *M* tumors, from equally many patients. We denote the *m*-th row of this matrix by *Y*_*m*_. This is a binary vector of length *N*, representing the mutation status of all genes (mutated/normal) in the *m*-th tumor with *Y*_*m,g*_ = 1 indicating that the gene *g* is mutated and *Y*_*m,g*_ = 0 otherwise. To allow reference to the status of only a subset of genes *S* ⊂ {1, … *N*} in the *m*-th tumor, we introduce a similar notation *Y*_*m,S*_. We use 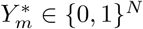 to denote a latent (noise free) gene status vector, where 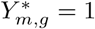 indicates that gene *g* is a driver mutation for the *m*-th sample. Finally, we model the biological noises that arise during DNA sequencing and data processing using two parameters *δ, ϵ* [0, 1]. Specifically, *E* denotes probability of a passenger mutation (i.e., false positive in the point of view of recovering the driver mutations) and *δ* denotes flip-back probability which models error sources such as dropout during sequencing (i.e., false negative).

### Probabilistic Generative Process

A graphical model representation of our generative probabilistic model is shown in Fig 2. We assume that given the pathway progression model, the tumors are independent. Hence, it suffices to describe the generative model for a single tumor *Y*_*m*_, *m* ∈ {1, …, *M*}.

**Fig 2.**
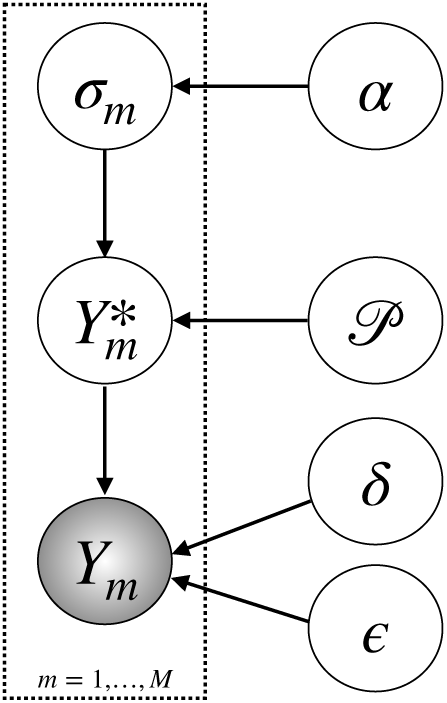
Generative process and the underlying graphical model for the observed mutation matrix.

#### Algorithm 2

Calculation of the likelihood *p*(*Y* | 𝒫 *α*; *ϵ*; *δ*)

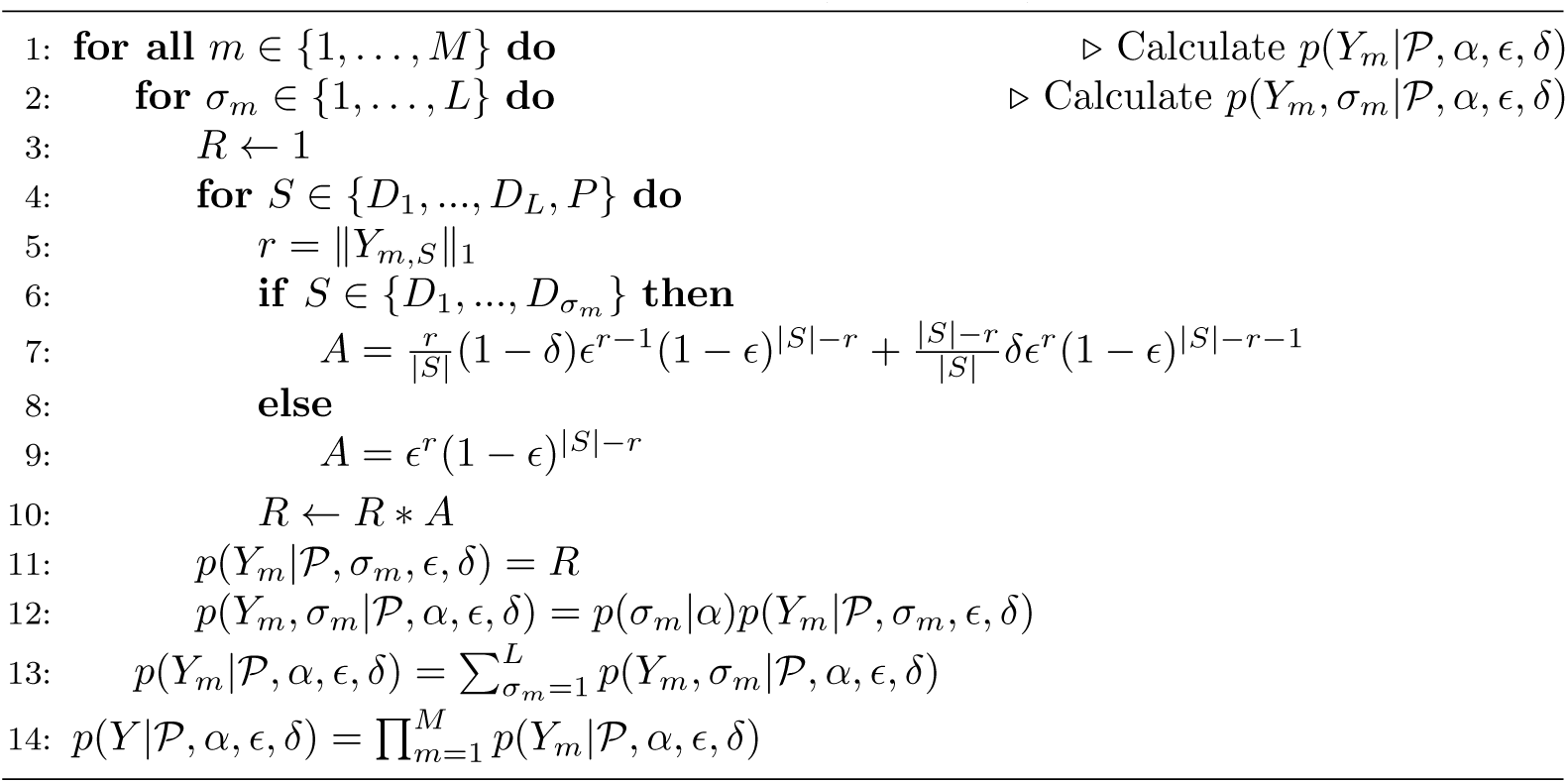

First, the latent progression stage *σ*_*m*_ ∈ {1, …, *L*} is sampled from a categorical distribution with parameter *α* = (*α*_1_, …, *α*_*L*_), i.e., *p*(*σ*_*m*_ = *l*) = *α*_*l*_ for *l* ∈ {1, …, *L*}. We use a fixed *α* = (1*/L*, …, 1*/L*). However, the parameter *α* can be arbitrarily chosen based on domain knowledge. We can even straightforwardly extend the graphical model to have a prior distribution on *α*, say a Dirichlet distribution, and infer the posterior belief on alpha given the data. For each *l* ∈ {1, …, *σ*_*m*_}, exactly one gene *g* ∈ *D*_*l*_ is mutated to construct 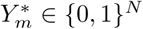. This procedure ensures mutual exclusivity of the genes: 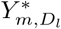 is a one-hot binary vector of dimension |*D*_*l*_| for *l* ∈ {1, …, *σ*_*m*_} and a zero vector everywhere else. We refer to the subsequent sub-process acting on 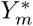 as corruption, since without this process, the driver mutations would be easily read off from the tumor. When generating the observed data *Y*_*m*_ given 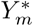, the ones may be flipped back to zero with probability *δ*. Alternatively, the zeros may turn into ones in our observed mutation status vectors with probability *ϵ*. The latter partly models technical problems such as mapping, misalignment and so on, but in particular that in a tumor any gene may acquire passenger mutations. In particular, even a so-called driver gene that belongs to a driver pathway in which another gene has been previously mutated may acquire a passenger mutation. Here, since the previous mutation has already affected the corresponding biological pathway, the mutation in the second gene does not confer any selective advantage to the tumor, and in this sense, the second mutation is a passenger mutation. The graphical model for the generative process can be found in Fig 2. The procedure for generating synthetic data, i.e., mutated tumor genomes based on our generative model, is also provided in Fig 2.

## Methods

We desire to estimate the posterior probability distribution for a given collection of tumors as well as to perform model selection, i.e., determine the number of driver pathways *L* by computing the marginal likelihood given *L*. An algorithm for computing the likelihood constitutes a crucial part of both these tasks.

**Fig 3.**
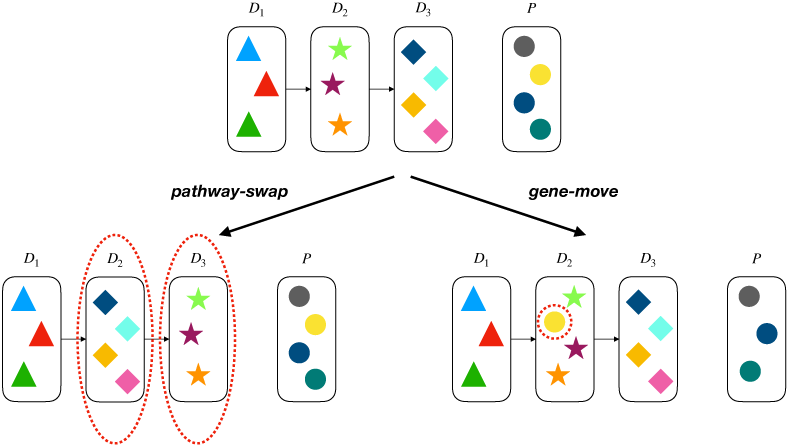
Two types of moves we use for progression model proposal.

### Computing the Likelihood

Let *Y* = {*Y*_*m*_ : *m* ∈ {1, …, *M*}} be a mutation matrix for a collection of tumors, 𝒫 = (*D*_1_, *D*_2_, …, *D*_*L*_, *P*) be a pathway progression model, and *α* be our prior distribution for the progression stages of the tumors. Denoting the bits in *Y*_*m*_ corresponding to pathway *S* by *Y*_*m,S*_, we can calculate *p*(*Y* | 𝒫, *α, ϵ, δ*) using Algorithm 2. The derivation steps of the algorithm and a faster implementation using look-up tables are presented in S1 Appendix.

### Markov Chain Monte Carlo algorithm to train the model

In this section, we describe an MCMC algorithm to generate samples from the posterior distribution of the latent quantities given the observations *Y* for a fixed model length *L*: *π*(𝒫, *ϵ, δ Y, L, α*). We apply Gibbs steps to sample the progression model and the error parameters iteratively for *T* MCMC iterations:

- Initialize 𝒫^0^, *ϵ*^0^, *δ*^0^.
- For *t* = 1,…, *T*:

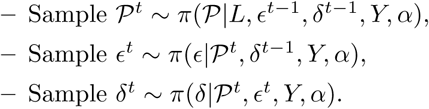

As sampling directly from the conditional posteriors is challenging, we use the Metropolis-Hastings algorithm to sample each of these latent variables, which renders our algorithm as a type of *Metropolis-within-Gibbs sampler* (see e.g., chapter 10.3.3 of [17]).

#### Sampling the progression model

Considering a uniform prior distribution for the progression structure given the model length *L* (in the space of all progression models of length *L*), we have:

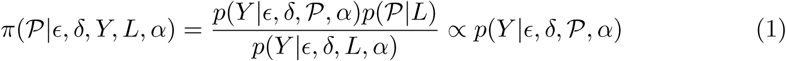

We use a Metropolis-Hasting sampler to generate progression model samples from this distribution. Our proposal function consists of two types of moves that we call *gene-move* and *pathway-swap*. We choose the type of move randomly using a Bernoulli distribution. For the case of *gene-move*, we select a gene uniformly at random and move it to a driver pathway or the set of passengers with a uniform distribution. For the case of *pathway-swap*, we select two driver pathways uniformly at random and swap their positions in the progression structure. According to our experiences, this type of moves can help the model to get out of locally optimal points, where the pathways are placed in a non-optimal order and the mutual exclusivity and the progression conditions are acting against each other, preventing the algorithm from moving towards more likely progression structures by moving one gene at a time. The acceptance ratio is given by,

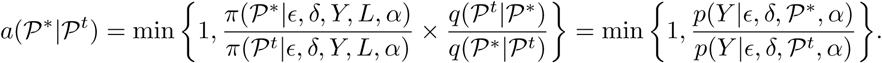

#### Sampling the error parameters

The conditional distribution for *ϵ* is given by,

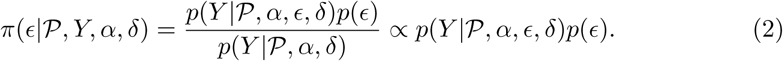

We use Gaussian random walk proposal, that is, we sample 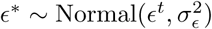. This leads to the following as the acceptance ratio,

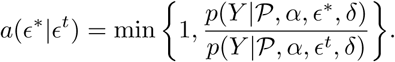

We use a similar Metropolis-Hasting sampler to sample our flip-back probability parameter *δ*. The variance parameters are chosen to be small, for example we use 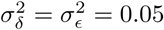 in our experiments. These variance parameters can also be adaptively selected as outlined in [18].

### Model selection

The MCMC sampler can be utilized for model selection. Model selection for pathway progression model involves finding a suitable value *L* for the number of pathways, called model length. We assume that we have a set of ℒ candidate model lengths, and that we are interested in computing the posterior probability of a model length *L* ∈ ℒ given the observation. Assuming a uniform prior on the model length, it suffices to compute the *model evidence p*(*Y* |*L*). One approach to estimate this quantity is to consider

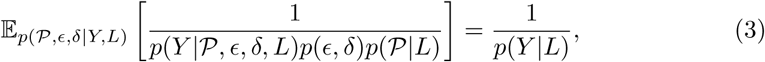

where the expectation can be estimated using the MCMC samples, leading to an estimate for *p*(*Y* | *L*). More details on the model selection procedure can be found in S2 Appendix.

## Synthetic Data Simulations

In this section, we use an extensive set of experiments on synthetic datasets to demonstrate the accuracy and efficiency of our method and, in particular, its superior performance compared to the earlier ILP-based approach [16]. For the synthetic data simulations, having the generative model 𝒫, we calculate a performance metric called *POCO* (Percentage Of Correct Ordering of genes). To this end, considering an inferred model 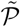, we go over all pairs of genes and for each pair, we check if their position with respect to each other (gene 1 before gene 2/gene 1 after gene 2/two genes in the same pathway) in 𝒫 is preserved in 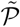. *POCO* is the percentage of the gene pairs with their relative position preserved.

### Experiment 1: Known driver genes

In this experiment, we have generated a set of synthetic datasets with fixed flip-back probability *δ* = 0.3, and various back-ground mutation rate *ϵ* and number of patients. We have distributed 25 genes in 5 driver pathways with 4 scenarios for the pathway sizes, i.e., number of genes in the pathways:

- *uniform*, where the sizes are (5, 5, 5, 5, 5),
- *increasing-by-2*, where the pathway sizes are (1, 3, 5, 7, 9)
- *decreasing-by-2*, where the pathway sizes are (9, 7, 5, 3, 1)
- *random*, where we randomly order the genes in a row and put 4 separating borders uniformly at random in 24 possible spots between genes.

Fig 4 shows the experiment results. As shown in this figure, while the ILP has difficulties with handling a large number of patients (leading to drop in its performance for large datasets due to inability to converge), our method effectively takes advantage of the statistical power arising from the increasing number of patients to improve its performance. This figure also shows the robust performance of our algorithm in all background mutation rates and pathway size scenarios compared to the ILP algorithm. Moreover, our MCMC samples can be used to estimate the error parameters. A comprehensive analysis of our error estimate performance can be found in S3 Appendix.

**Fig 4.**
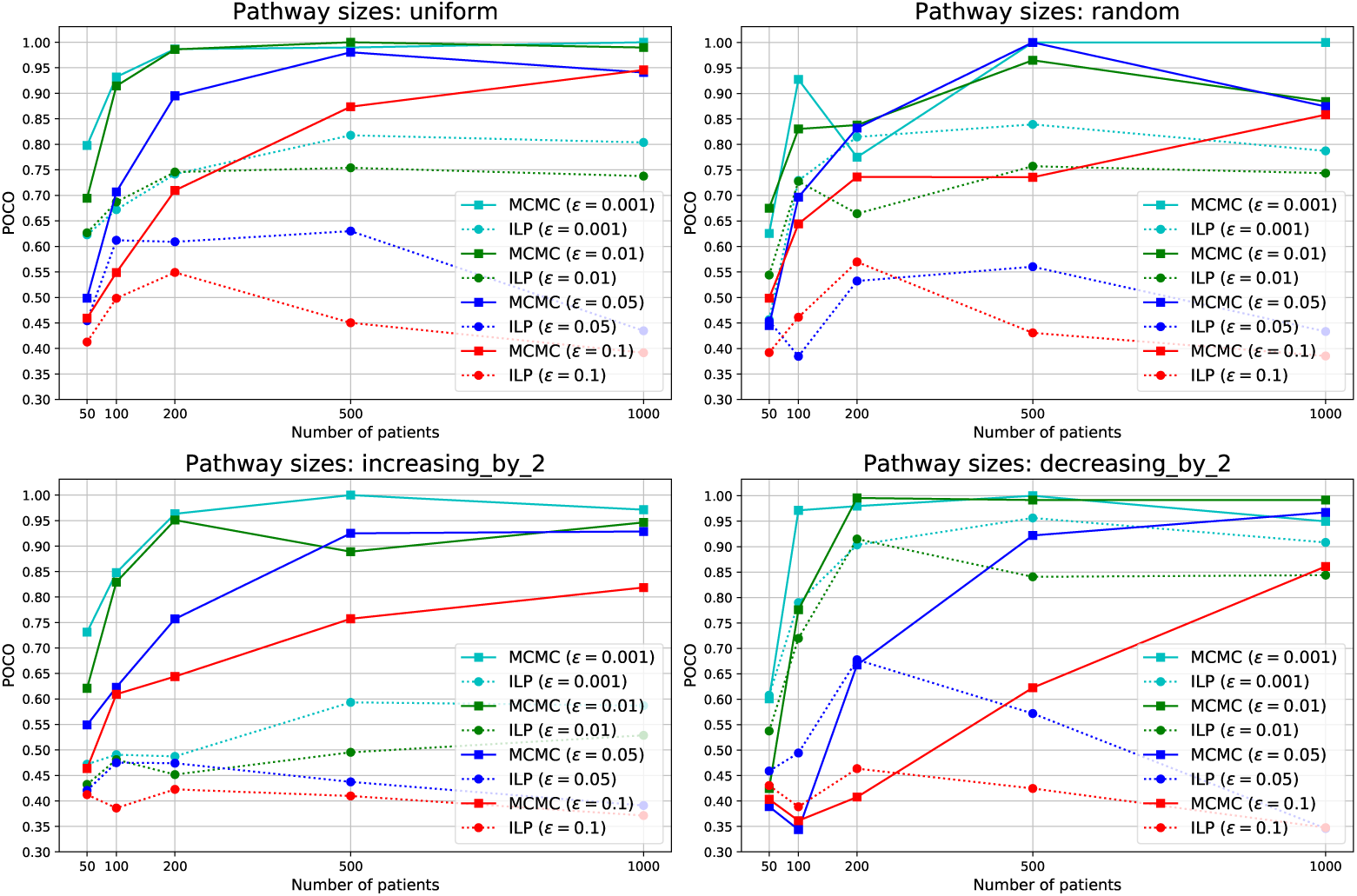
POCO, averaged over 10 datasets, for various *ϵ* values and pathway size scenarios. The flip-back probability is set to *δ* = 0.3 in all the datasets.

### Experiment 2: Unknown driver genes

In this experiment, we demonstrate the effect of adding passengers to the pool of genes. To this end, we generated datasets with 5, 25 and 100 passenger genes. For each case, we have constructed 10 datasets with 500 patients, flip-back probability of 0.3, background mutation rate of 0.05 and 25 genes uniformly distributed in 5 driver pathways.

While our MCMC algorithm does not require any information on the error parameter, the ILP algorithm requires the background mutation rate as an input to the algorithm, if we want to allow for assigning genes to the set of passengers. We have used 3 versions of the ILP algorithm with background mutation rates of 0.01, 0.05 and 0.1, to investigate the likely disruptive effect of incorrect background mutation rate input on the ILP method. We emphasize that the optimal background mutation rate parameter is not typically known in biological data. Hence, having a strong dependency on this parameter is a significant drawback for the ILP-based method.

In Table 1, we have shown the performance of the competitors using POCO measure, driver detection F1 score, and specific pathway detection F1 score. As shown in this table, the ILP given the true generative background mutation rate of 0.05 performs best among the ILPs, as expected. However, our algorithm significantly outperforms all the ILP competitors, even the one with the extra knowledge of the true generative error parameter. More detailed results on the detection of the genes in specific pathways can be found in S4 Appendix.

**Table 1.** POCO, F1 scores for driver detection and F1 scores for specific pathway detection (averaged over pathway 1 to 5) in Experiment 2.

| Method | POCO |  |  | F1 score (driver detection) |  |  | F1 score (pathway detection) |  |  |
| --- | --- | --- | --- | --- | --- | --- | --- | --- | --- |
|  | 5 passengers | 25 passengers | 100 passengers | 5 passengers | 25 passengers | 100 passengers | 5 passengers | 25 passengers | 100 passengers |
| MCMC | 0.942 | 0.936 | 0.951 | 0.974 | 0.960 | 0.935 | 0.916 | 0.877 | 0.868 |
| ILP($\epsilon = 0.01$) | 0.568 | 0.423 | 0.285 | 0.909 | 0.667 | 0.343 | 0.392 | 0.238 | 0.086 |
| ILP($\epsilon = 0.05$) | 0.772 | 0.808 | 0.856 | 0.882 | 0.864 | 0.779 | 0.389 | 0.379 | 0.304 |
| ILP($\epsilon = 0.1$) | 0.551 | 0.494 | 0.713 | 0.493 | 0.428 | 0.400 | 0.380 | 0.333 | 0.276 |

### Experiment 3: Model length selection

In this experiment, we demonstrate our model length selection performance. To this end, we have generated datasets using model length parameters from 2 to 9 with 50, 100, 200, 500 and 1000 patients, 25 driver genes uniformly distributed in the driver pathways, 175 passenger genes, background mutation rate of 0.01, and flip-back probability of 0.3. We have constructed 10 datasets using each model length and used our method with model length candidates from 2 to 9.

Fig 5 shows the confusion matrices for various numbers of patients. As shown in this figure our performance gets better as the number of patients in the dataset increases. However, we emphasize that as shown in the POCO tables in the second row of Fig 5, even for the cases that we have not correctly identified the model length, the genes ordering with respect to each other is recovered up to a significant level, which is of great importance. Consider the case of 100 patients, for instance. We can see from the corresponding confusion matrix in the first row of Fig 5 that the element (6, 2) equals 4. This means that in 4 simulations with the generative model length of 6, our algorithm has mistakenly inferred the model length equal to 2. However, looking at the corresponding element in the averaged poco matrix (in the second row of Fig 5), we see that in these 4 simulations, our inferred model has achieved a POCO score of 0.88 in average, which seems a quite satisfying result for such a small number of patients.

**Fig 5.**
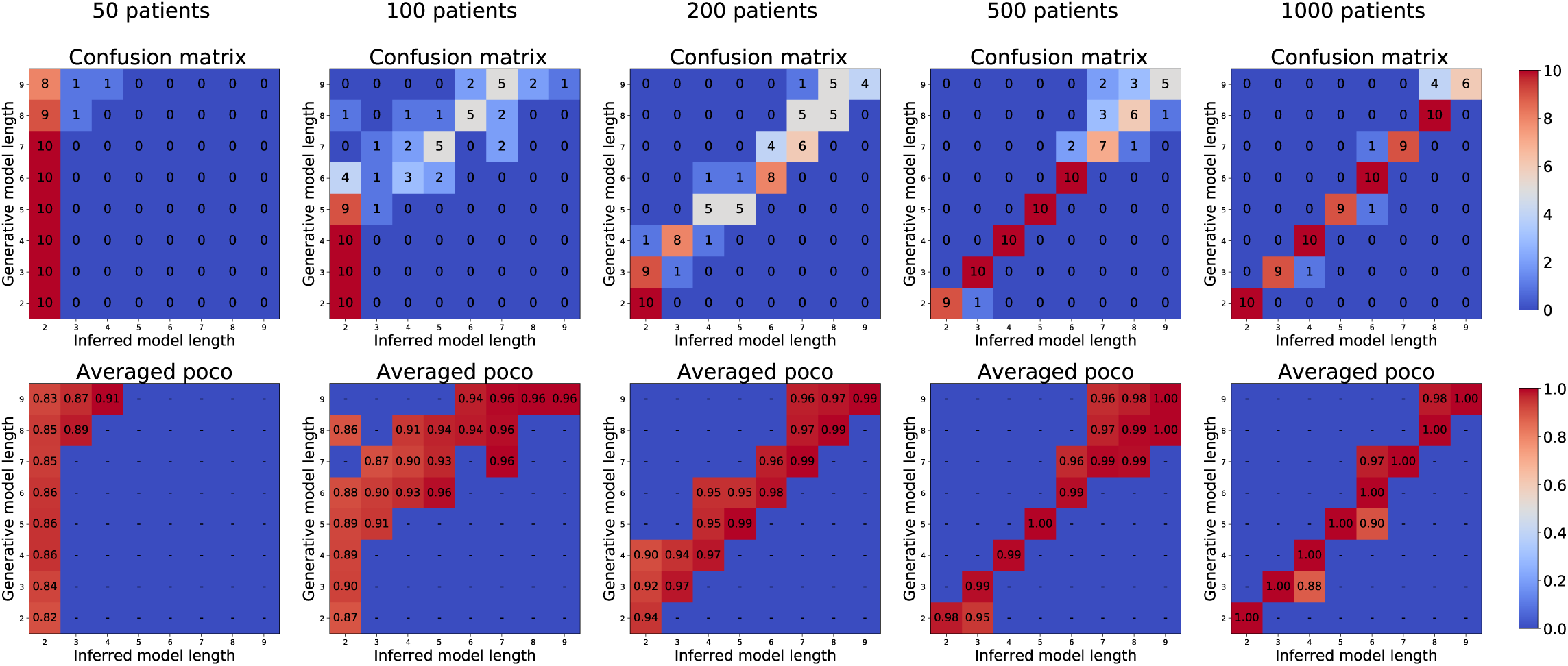
Confusion matrices and averaged POCO scores of our MCMC algorithm in Experiment 3. The matrices in the first row show the confusion matrices for simulations with various number of patients. The element (*i, j*) in each one of the matrices in the first row shows the number of experiments in which the generative model length was *i* and the inferred model length was *j*. The matrices in the second row show the averaged POCO scores of the inferred models in each scenario. Here, the element (*i, j*) shows the average POCO score for the cases in which the generative model length was *i* and the inferred model length was *j*.

## Biological Data Analysis

We analyzed two large biological datasets of colorectal adenocarcinoma (COADREAD) and glioblastoma multiforme (GBM) from IntOGen-mutations [19] and compared our algorithm against our implementation of the ILP-based method in [16]. To this end, we first filtered out all the silent mutations as a preprocessing step. Alongside the datasets, IntoGen has published a list of potential driver genes for each cancer type. We call the genes in these lists the *potential driver genes* and use only those genes in our input matrices for both our algorithm and the ILP-based competitor. These input matrices can be found in Fig 6-A and Fig 7-A. In these matrices, the rows and columns represent the genes and the tumors, respectively. A black rectangle in position (*i, j*) means that tumor *j* has a mutation in gene *i*. After the preprocessing steps, we have 9465 genes and 290 patients with 24 genes in the list of potential driver genes for GBM, and 9169 genes and 193 patients with 37 potential driver genes for COADREAD.

**Fig 6.**
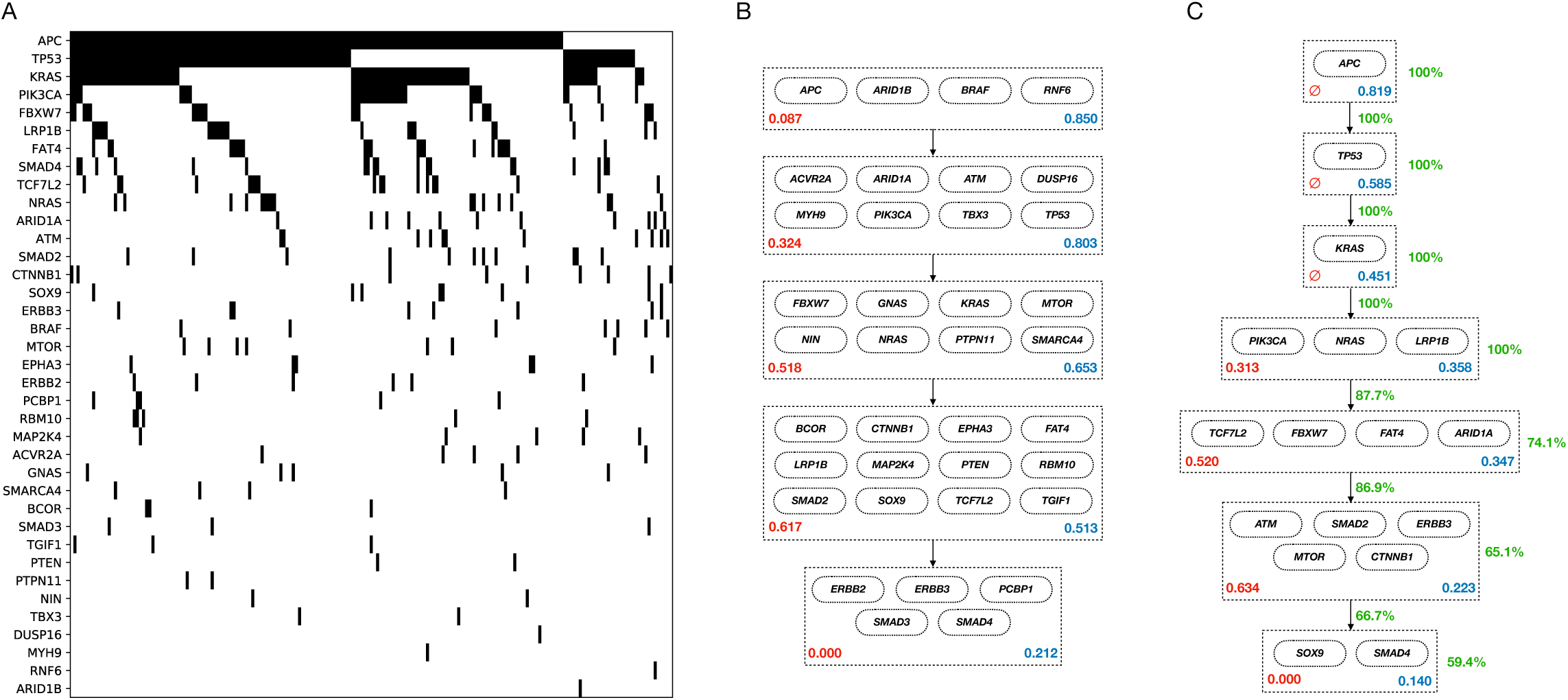
The COADREAD dataset analysis. A: The dataset representation. The genes are sorted based on their mutation frequencies. B: ILP-based method inferred model. C: Our MCMC method inferred model.

**Fig 7.**
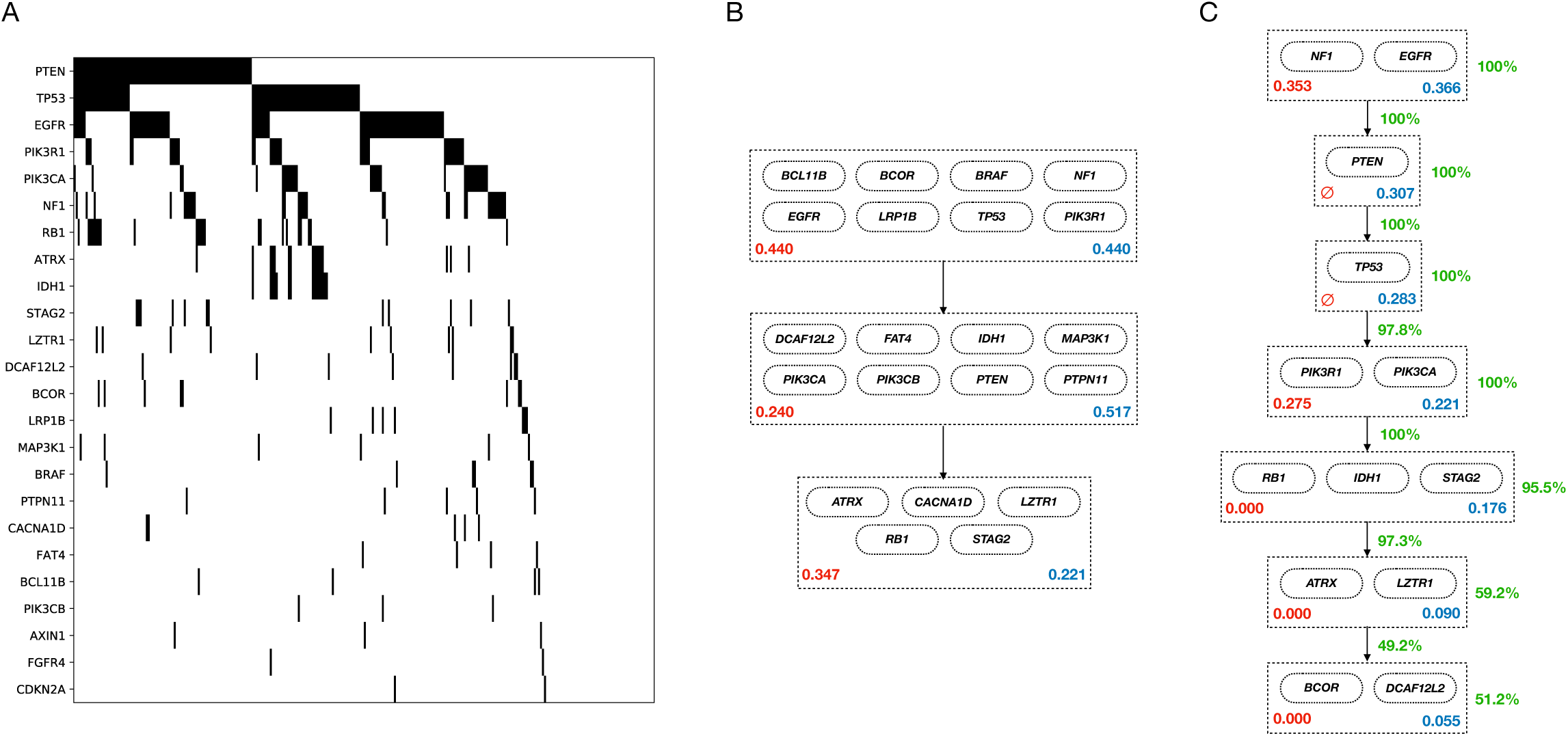
The GBM dataset analysis. A: The dataset representation. The genes are sorted based on their mutation frequencies. B: ILP-based method inferred model. C: Our MCMC method inferred model.

For each dataset, we performed model selection routines with candidate lengths from 2 to 30 and chose the best model length. The output models for both our algorithm and the ILP-based method are shown in Fig 6 and Fig 7. The MCMC output models shown in these figures are consensus-like models, constructed using our samples from the posterior distributions of the progression models. A pseudo-code description of the algorithm we used to construct our consensus-like output models is provided in S6 Appendix. This algorithm takes the MCMC model samples as input and outputs a progression model that can be considered as *the average model*. We emphasize that in our analyses, the consensus-like models were pretty close to the MCMC samples with the highest likelihoods in both datasets.

After we constructed the average model, we calculate our confidence on the pathways and their respective position in the progression model as follows. For each pathway, we go over all pairs of genes in the pathway. Our confidence in the pathway is the averaged percentage of our MCMC samples with these pairs of genes in the same pathway. To calculate our confidence on an arrow connecting pathway 1 to pathway 2, we go over all pairs of genes (gene_1_, gene_2_) with gene_1_ in pathway 1 and gene_2_ in pathway 2. Our confidence on the arrow is the percentage of our MCMC samples that have put gene_1_ before gene_2_. We have shown these confidence metrics using green numbers beside the arrows and pathways in the depicted models.

To assess the performance of our algorithm against the ILP-based competitor, we define a metric named ME (Mutual Exclusivity) score for each pair of genes, denoted by *S*_ME_:

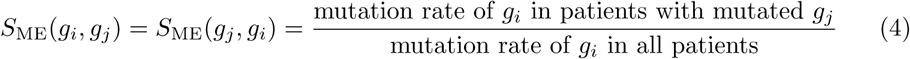

As the average ME score of the pairs of genes in the identified pathways is smaller, we have a higher level of mutual exclusivity in the pathway, and hence a better model. We have shown the ME scores of the pathways identified by both algorithms using red numbers inside the depicted pathways in Fig 6 and Fig 7. Moreover, the fraction of the tumors having at least one mutation in each pathway is shown using the blue numbers inside the pathways. The first driver pathway having the highest fraction of mutations and a decreasing trend afterward implies that the progression condition holds in the inferred model.

As shown in Fig 6, our algorithm results in a progression model of length 7 for the COADREAD dataset. The first pathway includes *APC*, a tumor suppressor gene. Mutation and inactivation of this gene is known to be an early event playing a key role in colorectal cancer tumorigenesis [20]. The second pathway includes *TP53*, another tumor suppressor, which is highly mutated in colorectal cancer. Mutant p53 is also shown to have an oncogenetic role in colorectal cancer through gain-of-function mechanisms [21]. The third pathway includes *KRAS*. Mutated *KRAS* is known to be highly associated with colorectal cancer [22]. *KRAS* mutation in colorectal cancer is also known to be a subsequent event after a mutation in *APC* [23], a temporal order of mutations that is recovered by our algorithm as well.

As shown in Fig 7, our algorithm inferred a progression model of length 7 for the GBM dataset. The first pathway includes *EGFR* and *NF1*, two genes known to be the main drivers of the *classical* [24] and *Mesenchymal* [25] subtypes, respectively. *PTEN* in the second pathway is a well-known tumor suppressor known to be involved in regulation of glioblastoma oncogenesis [26]. The third pathway includes *TP53*, another well-known tumor suppressor gene, which is known to play an important role in various cancer types. In glioblastoma in particular, the *p53-ARF-MDM2* pathway is reported to be deregulated in 84% of the patients and 94% of the cell lines [27]. The fourth pathway includes two class IA PI3K subunit genes *PIK3CA* and *PIK3R1*. Mutations in *PI3K* survival cascade is known to be highly associated with glioblastoma [28]. The genes in the fifth pathway *IDH1, RB1* and *STAG2* are also known to be associated with the cancer progression in brain glioblastoma [29], [30], [31].

Comparing our results against the ILP-based method, we see that the ILP-based method puts a lot of genes with low mutation rates in the driver pathways, while our algorithm keeps its focus on the highly mutated genes. This happens since the ILP is trying to minimize the number of bits that are needed to get flipped to make the dataset perfectly following the error-less linear progression model. Hence, the ILP tends to put some not necessarily driver genes with low mutation rates into the driver pathways, as it does not affect the ILP cost that much. As an extreme example, if a gene is not mutated in the dataset, the model cost does not depend on where the ILP puts the gene. Our algorithm on the other hand implicitly considers equal importance for the genes in a pathway, which prevents us from putting less highly mutated genes in the driver pathways. Moreover, while our MCMC samples provide us with proper confidence metrics on the inferred models, the ILP-based method has to use bootstrapping techniques and re-run the algorithm over and over to provide some confidence metrics.

## Discussion

We investigated the progression patterns in cancer. To this end, we developed a probabilistic model that tries to capture the patterns of progression and mutual exclusivity among the genes involved in cancer. We designed an efficient MCMC algorithm to make inferences on the progression model. We demonstrated the superior performance of our algorithm compared to a previously introduced ILP-based method on a wide set of synthetic data simulations. We also analyzed two biological datasets on colorectal cancer and glioblastoma and showed that our inferred progression models can be better validated compared to the models suggested by the ILP-based counterpart.

## Supporting Information

## S1 Appendix. Likelihood calculation details

We want to calculate *p*(*Y* |*𝒫, σ*_1:*M*_, *ϵ, δ*), where *Y* = {*Y*_*m*_ : *m* ∈ {1, …, *M*}} is our observation matrix, *σ*_1:*M*_ = (*σ*_1_, …, *σ*_*M*_) is the given vector of progression stages of the tumors, and 𝒫 = (*D*_1_, *D*_2_, …, *D*_*L*_, *P*) is our pathway progression model. Since the tumors are independent given 𝒫 and *σ*_1:*M*_,

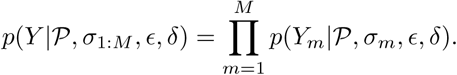

We separate the bits corresponding to different pathways as:

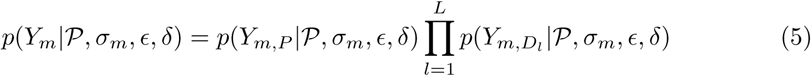

In order to calculate each term of form *p*(*Y*_*m,S*_|*𝒫, σ*_*m*_, *ϵ, δ*) in (5), we marginalize over all possible noise-free vectors 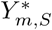:

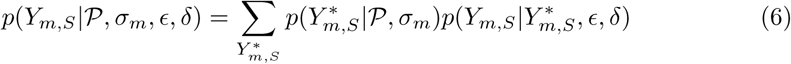

If 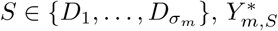 has to be a one-hot binary vector of length |*S*|. Denoting the number of ones in the observed *Y*_*m,S*_ by *r*, we have:

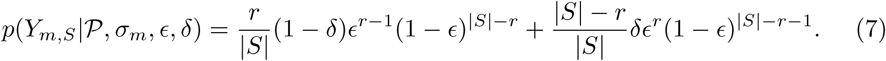

The first summand in this expression corresponds to the probability of the 1 in the latent 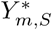 being among our *r* observed 1’s in *Y*_*m,S*_ (which is the case with probability of *r/*|*S*|), times the probability of getting to *Y*_*m,S*_ from such a 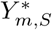. In this case, *Y*_*m,S*_ is obtained by the 1 in 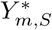 being kept from flip-back, followed by passenger mutations in *r* − 1 genes (leading to the total of *r* observed mutations) and no false positives in the remaining |*S*| − *r* genes. Similarly, the second summand in (7) corresponds to the probability of the 1 in the latent 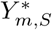 being among our |*S*| − *r* observed 0’s in *Y*_*m,S*_ (due to a flip-back). This is the case with probability of (|*S*| − *r*)*/*|*S*|, and if it is, then *Y*_*m,S*_ is obtained by a flip-back, followed by passenger mutations in *r* genes and no false positives in the remaining |*S*| − *r* − 1 genes.

If 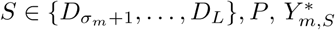 has to be a vector of |*S*| zeros. Hence, observing *r* ones in *Y*_*m,S*_, we have exactly *r* false positives, leading to:

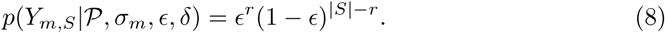

### Unknown progression stages

In practice, we do not have the progression stages of individual tumors. Fortunately, we can marginalize out the progression stages vector *σ*_1:*M*_ using the independence assumption over the samples given the pathways:

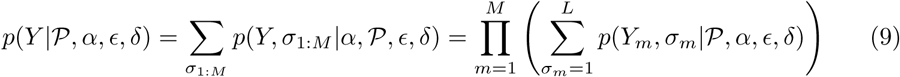

In order to calculate *p*(*Y*_*m*_, *σ*_*m*_|*𝒫, α, ϵ, δ*), we can write it as

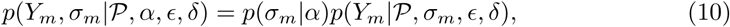

where the first term is the prior belief on the stages being *σ*_*m*_ and the second term is given by (5). We consider a uniform prior on the progression stages *σ*_1:*M*_, i.e., *α* = (1*/L*, …, 1*/L*). However, an alternative prior can be chosen.

In the following subsection, we describe a systematic likelihood calculation scheme, which can prevent us from repetitive calculations while going over our *M* tumors in the data.

### Fast Likelihood Calculation

Given a progression model 𝒫 = (*D*_1_, …, *D*_*L*_), and the data matrix *Y*, we form a matrix *C* of shape (*M, L*), where *C*_*i,j*_ is the number of mutations of tumor *i* in driver pathway *j*. Denoting the size of our largest pathway by *z* = max_*l*∈[*L*]_ |*D*_*l*_|, we form two look-up tables in form of zero matrices 𝒜 and ℬ of shape (*z, z*). We modify algorithm 2 to check the lookup tables before repetitive calculations in lines 7 and 9. The modified procedure is provided in Algorithm 3.

## S2 Appendix. Model selection details

The posterior probability of a model length *L* ∈ ℒ given the observation can be written as

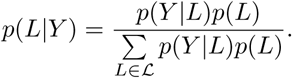

Our uniform prior on the model length corresponds to setting *p*(*L*) = 1*/*|ℒ |, hence, *p*(*L*|*Y*) ∝ *p*(*Y* |*L*). The equation in (3) can be simply derived as follows,

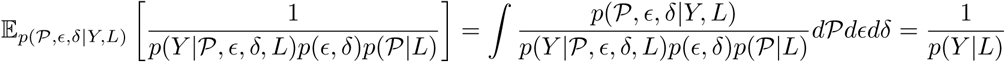

The MCMC estimate of the LHS in this equation will be

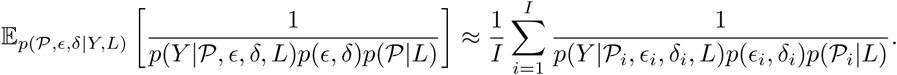

One potential problem is in the computation of *p*(𝒫_*i*_|*L*), where we need to calculate the cardinality of the space of valid pathway progression models of length *L*: |*𝒳* (*L*)|. Although enumeration over *𝒳* (*L*) appears intractable, there is a closed-form formula for computing its cardinality, |*𝒳* (*L*)|. Given a set of *N* genes, a valid progression of length *L* consists of *L* non-empty driver pathways and a set of passenger genes. Let *f*_*N*_ (*L*) denote the number of valid ways to allocate *N* genes to *L* non-empty driver pathways and the set of passengers (which can remain empty). We can calculate *f*_*N*_ (*L*) using the recursive formula

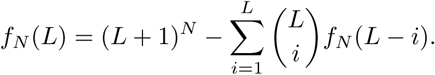

Note that *f*_*N*_ (*L*) is the total number of possible assignments, (*L* + 1)^*N*^, minus the number of invalid assignments. We count the number of invalid assignments with *i* empty driver pathways separately. There exist 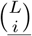 different choices for a set of *i* driver pathways to keep empty. For each case, we can have *f*_*N*_ (*L* − *i*) different valid assignment of genes to the remaining pathways, ensuring that none of the *L* − *i* remaining driver pathways are empty.

### Algorithm 3

Fast calculation of the likelihood *p*(*Y* | 𝒫, *α ϵ δ*)

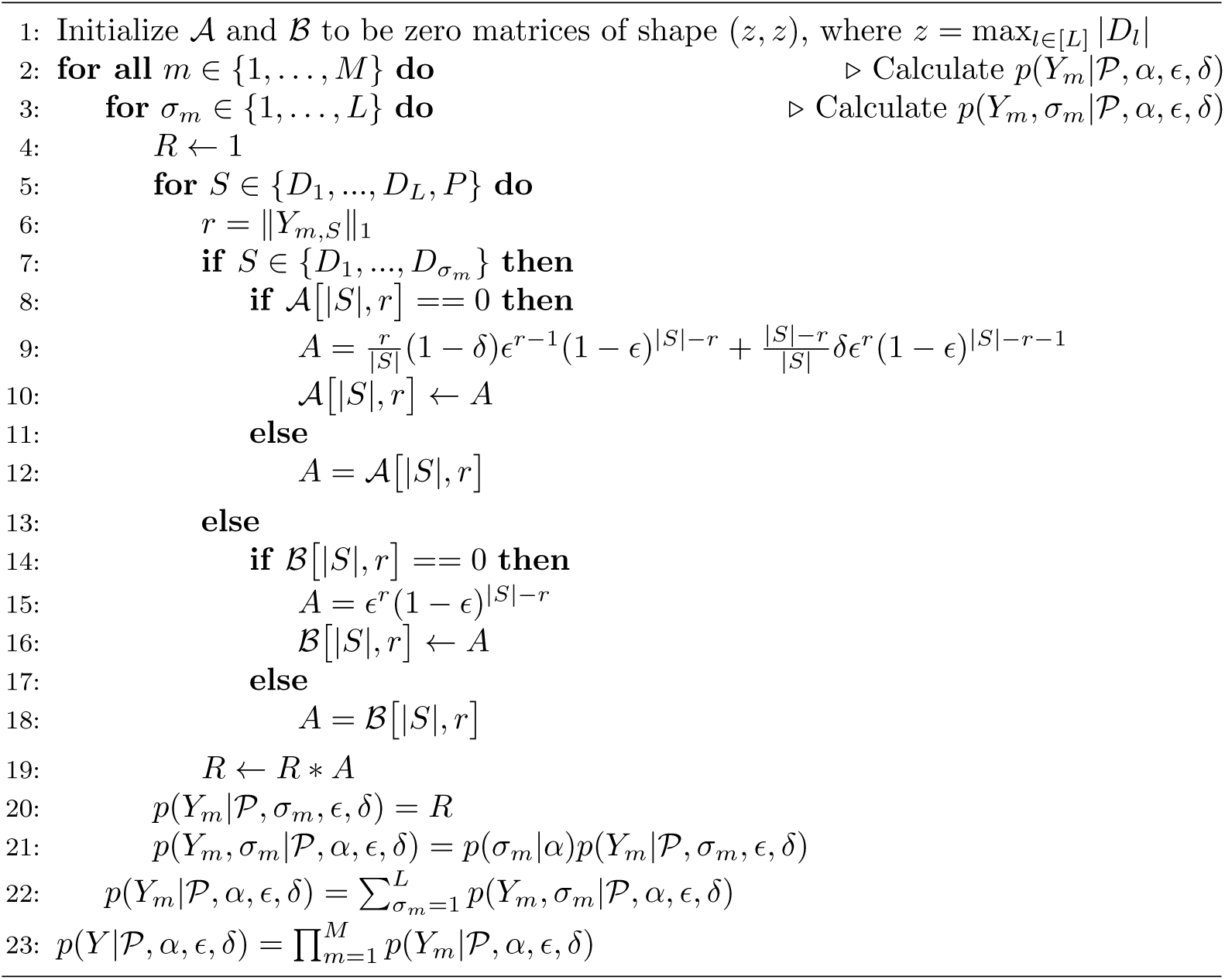

## S3 Appendix. Experiment 1 details

In all the experiments in this section, we have given a 60 seconds time limit to ILP. We have used MCMC with 2500 iterations (first 500 discarded as burn-in phase) with 0.9 probability of *gene-move* and 0.1 probability of *pathway-swap*, 10 Metropolis-Hasting iterations within each for error rate (*ϵ* or *δ*) update, and Gaussian random walk proposal for *ϵ* and *δ* with standard deviation of 0.05. All MCMC run times were less than 30 seconds.

One of the main advantages of a Bayesian approach such as ours, compared to the ILP algorithm, is the posterior distribution we can generate for the error parameters. Fig 8 shows our performance on estimations of the error parameters. The true (generative) values of the error parameters are shown using the red dotted lines. As shown in this figure, the estimates get more and more precise as the number of patients increases.

**Fig 8.**
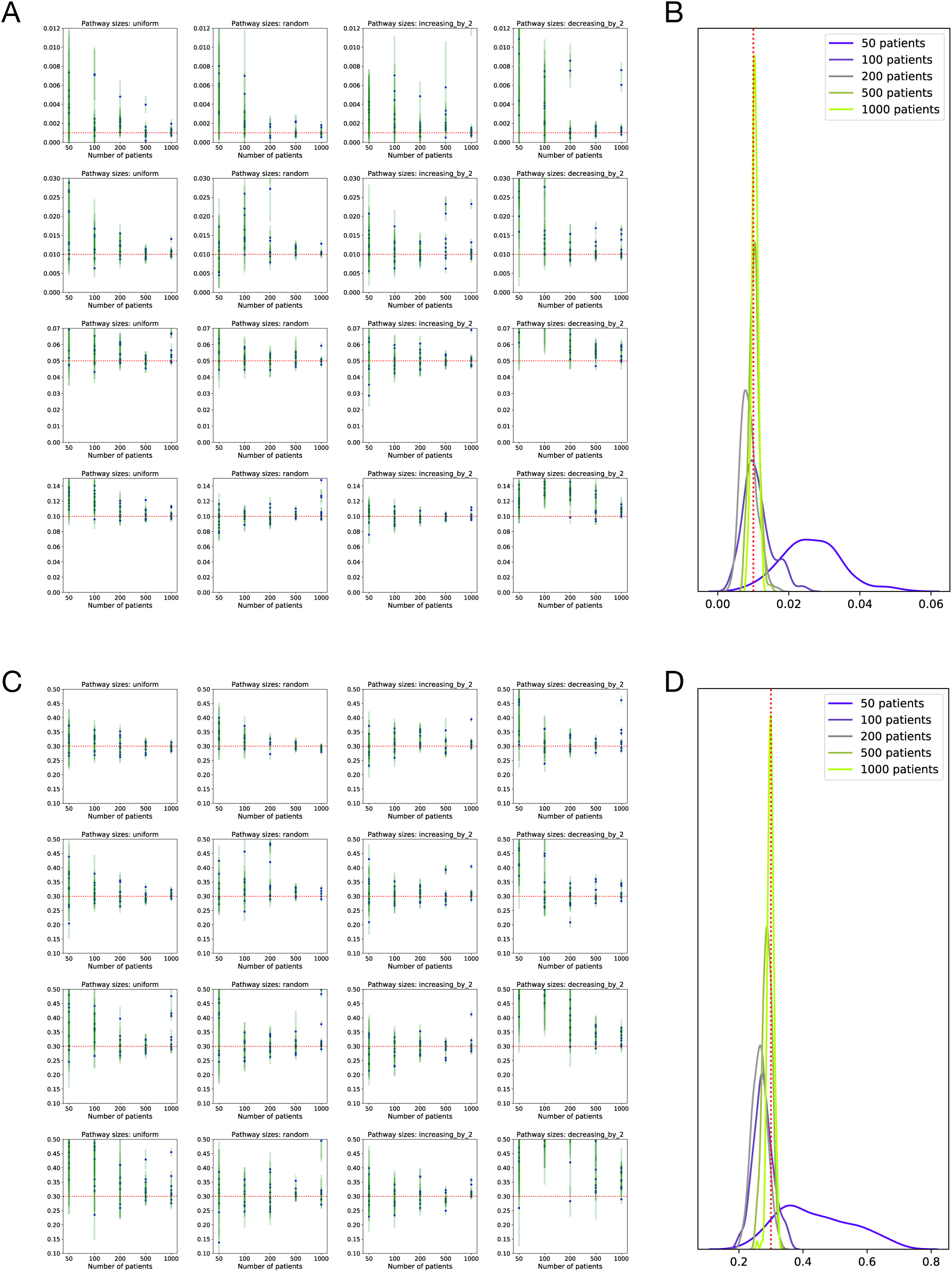
The estimated error parameters for Experiment 1. The red dotted lines show the true generative value for the error parameters. A: The posterior means (blue dots) and standard deviations (green lines) of the background mutation rates for all MCMC chains. B: Example KDEs for the background mutation rates. C: The posterior means (blue dots) and standard deviations (green lines) of the flip-back probabilities for all MCMC chains. D: Example KDEs for the flip-back probabilities.

## S4 Appendix. Experiment 2 details

In this experiment, for the MCMC algorithm, we have used the same setting as the previous experiment except for a prior passenger probability of 0.99 (for all genes) and an increment in the number of iterations to 10000 (first 2000 are discarded as burn-in phase). Each ILP run had a time limit of 600 seconds, while the MCMC run times in this setting take less than 60 seconds typically (one-tenth of our ILP time limit).

In Fig 9, we show the precision, recall and F1 score of the algorithms for the task of specific pathway identification. It can be seen that the detection of the genes in the last pathways is a harder task for all the methods, as expected. Our MCMC method outperforms all the ILP algorithms, even the one with the extra knowledge on the generative error parameter (ILP with *ϵ* = 0.05).

**Fig 9.**
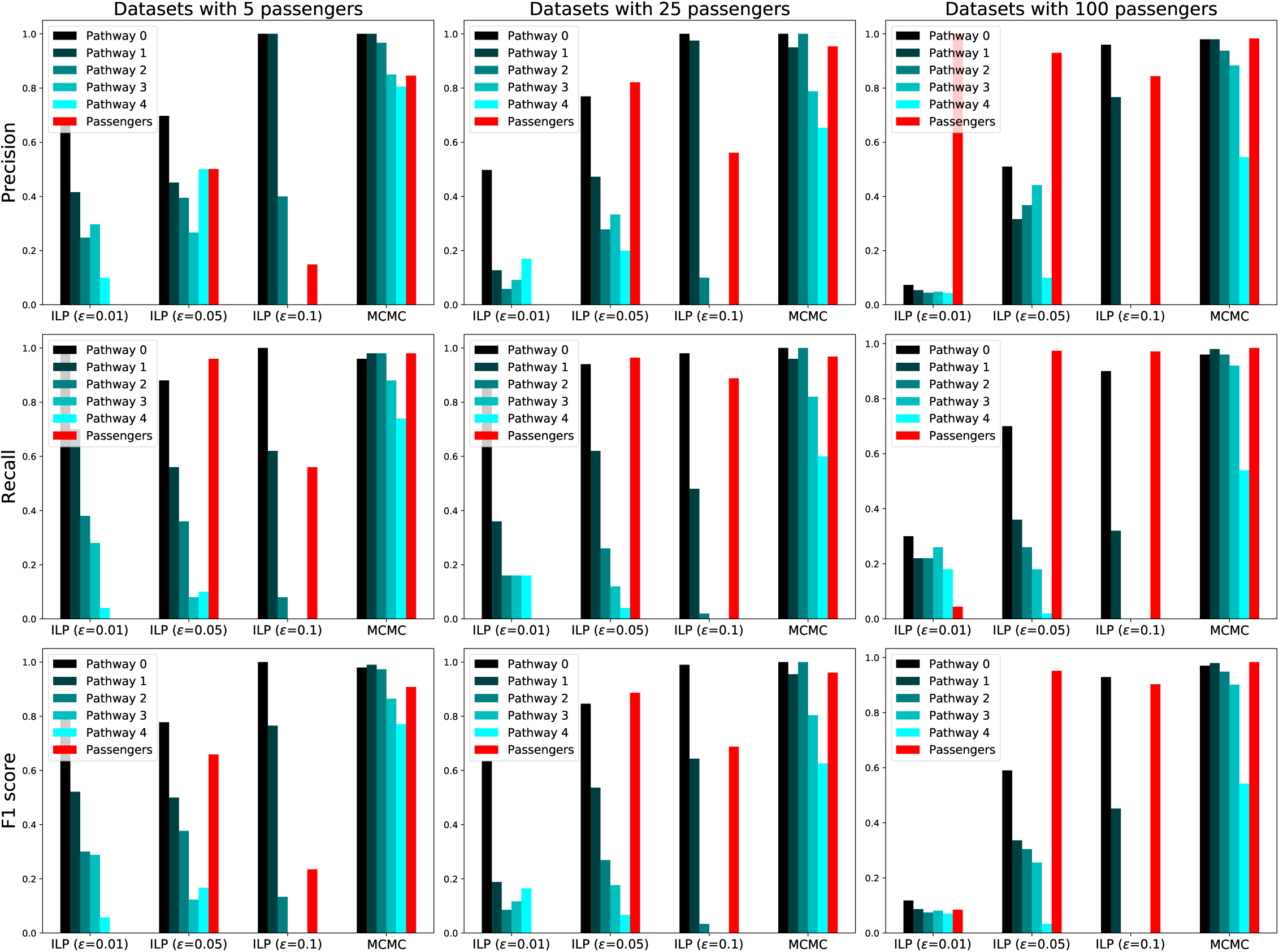
Precision, recall and F1 scores in detection of the genes in pathways 1 to 5 and the set of passengers in Experiment 2.

## S5 Appendix. Experiment 3 details

In this experiment, the MCMC parameters are exactly similar to the previous experiment, except for the varying model length. After running MCMC with various model length, we have chosen the inferred model length based on the dataset evidence provided by the MCMC algorithm.

## S6 Appendix. Averaging the MCMC model samples

We use Algorithm 4 to produce a consensus-like model from a set of models collected from the MCMC run. The goal of this algorithm is to output a progression model (of the same length as the input models) that preserves the genes respective orders as much as possible.

## S7 Appendix. Biological data analysis details

As the set of potential drivers typically includes less than one percent of the genes, the background mutation rate has to be roughly equal to the mutation rate of the dataset. We used this information to set the cost coefficient of the passenger genes in the ILP-based method and set our algorithm’s background mutation rate to be equal to the mutation rate of the dataset.

For our algorithm, we had 10000 MCMC iterations for each model length. We had 10 MH iterations for *δ* parameter within each Gibbs iteration. After running MCMC with various model lengths, we chose the inferred model length based on the dataset evidence provided by the MCMC algorithm. We used a thinning interval of 10 samples to collect less correlated samples and considered the last 500 samples to compute the consensus-like model as our output.

For the ILP-based method, we considered a time limit of 600 seconds for each ILP run. The cost coefficient of the passenger genes was set using the information on the background mutation rate. The model selection was performed using the procedure suggested by the corresponding publication on the same set of model length candidates as the one used for the MCMC (from length 2 to 30).

## Acknowledgments

This work was funded by ITN-CONTRA EU grant H2020 MSCA-ITN-2017-766030. This work was supported by the grant BD15-0043 from the Swedish Foundation for Strategic Research.

## Algorithm 4

Mixing MCMC samples from the posterior distribution of the progression models.

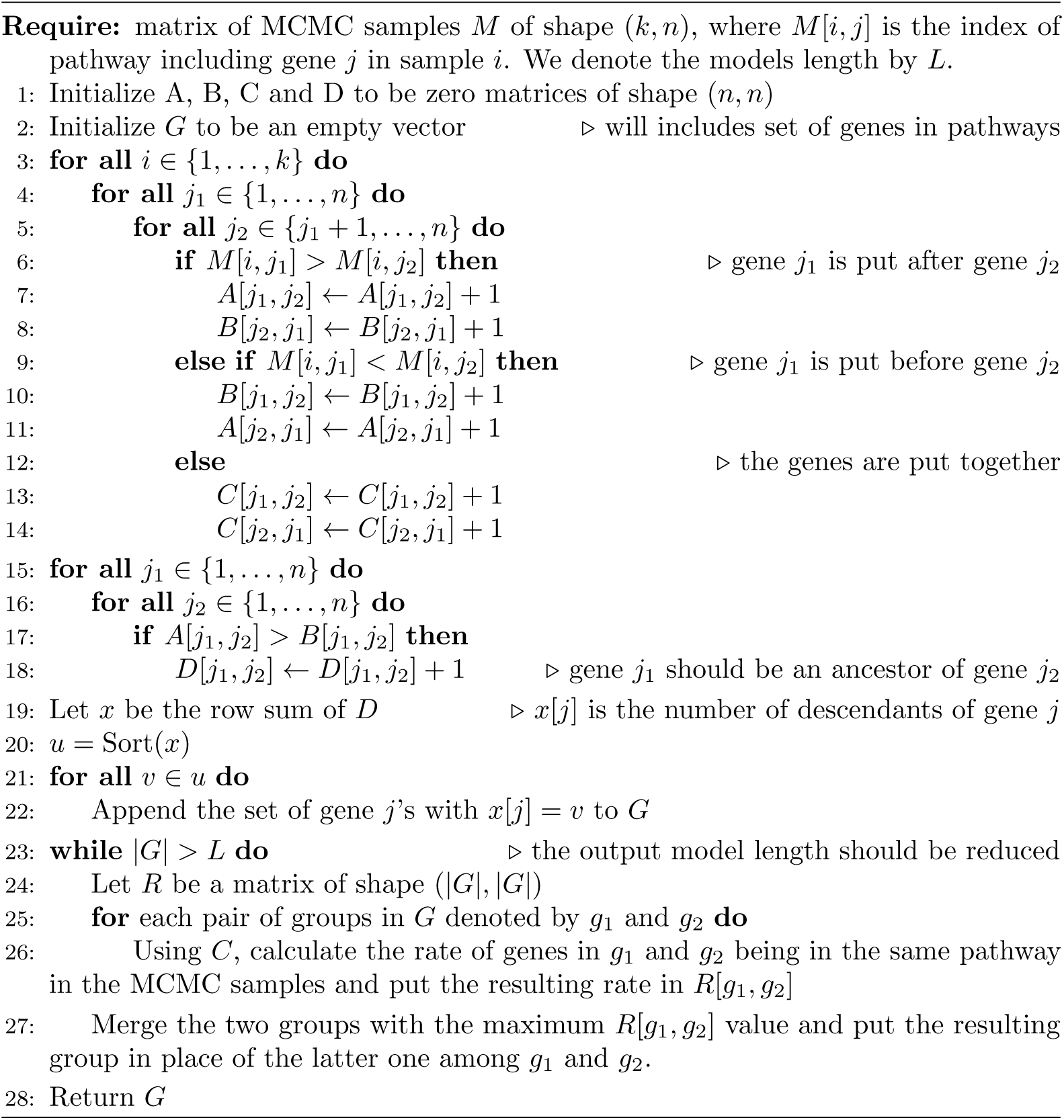

